## Supplemental Table 1 for "Bud23 promotes the progression of the Small Subunit Processome to the pre-40S ribosome in *Saccharomyces cerevisiae*"

| **Gene** | **Mutation** | **Reference** |
| --- | --- | --- |
| ***IMP4*** | D74G^1^ | This study. |
| ***IMP4*** | Y77C^1^ | This study. |
| ***IMP4*** | T92I^1^ | This study. |
| ***IMP4*** | S93T^1^ | This study. |
| ***IMP4*** | R94L^1^ | This study. |
| ***IMP4*** | R94C^1^ | This study. |
| ***IMP4*** | R94S^1^ | This study. |
| ***IMP4*** | R99H^1^ | This study. |
| ***IMP4*** | R99L^1^ | This study. |
| ***IMP4*** | S101W^1^ | This study. |
| ***IMP4*** | R116M^1^ | This study. |
| ***IMP4*** | N118D^1^ | This study. |
| ***IMP4*** | N118K^1^ | This study. |
| ***IMP4*** | N121I^1^ | This study. |
| ***IMP4*** | H143D^1^ | This study. |
| ***IMP4*** | R146G^1^ | This study. |
| ***IMP4*** | H156D^1^ | This study. |
| ***IMP4*** | H159R^1^ | This study. |
| ***IMP4*** | V170F^1^ | This study. |
| ***IMP4*** | H208D^1^ | This study. |
| ***IMP4*** | P252L^1^ | This study. |
| ***UTP2*** | A2D^1,2^ | (1) |
| ***UTP2*** | L6P^2^ | This study. |
| ***UTP2*** | K7E^2^ | This study. |
| ***UTP2*** | L9S^1^ | This study. |
| ***UTP2*** | F58S^2^ | This study. |
| ***UTP2*** | L148S^2^ | This study. |
| ***UTP2*** | F149S^2^ | This study. |
| ***UTP2*** | L151H^2^ | This study. |
| ***RPS28A*** | G24D^1^ | This study. |
| ***BMS1*** | D124Y^1^ | This study. |
| ***BMS1*** | G813S^1^ | This study. |
| ***BMS1*** | D843V^1^ | This study. |
| ***BMS1*** | A903P^1^ | This study. |
| ***BMS1*** | S1020L^1^ | This study. |
| ***UTP14*** | V754G^2^ | (2) |
| ***UTP14*** | I755T^1,2^ | (2) |
| ***UTP14*** | E757G^2^ | (2) |
| ***UTP14*** | A758G^1,2^ | (1) |
| ***UTP14*** | A760P^2^ | (2) |
| ***UTP14*** | W791L^1^ | This study. |
| ***UTP14*** | W794L^1^ | This study. |
| ***DHR1*** | R13G^1^ | This study. |
| ***DHR1*** | E360K^2^ | (3) |
| ***DHR1*** | E397D^1^ | This study. |
| ***DHR1*** | E402G^2^ | (3) |
| ***DHR1*** | D408Y^1^ | This study. |
| ***DHR1*** | E430K^1^ | (3) |
| ***DHR1*** | G432R^1^ | This study. |
| ***DHR1*** | G434D^1^ | This study. |
| ***DHR1*** | S511Y^1^ | This study. |
| ***DHR1*** | R563M^1^ | This study. |
| ***DHR1*** | D566Y^1^ | This study. |
| ***DHR1*** | F567L^2^ | (3) |
| ***DHR1*** | H593Y^2^ | (3) |
| ***DHR1*** | F594L^1^ | This study. |
| ***DHR1*** | R596C^2^ | (3) |
| ***DHR1*** | R596G^1^ | This study. |
| ***DHR1*** | R596S^1^ | This study. |
| ***DHR1*** | M744T^2^ | (3) |
| ***DHR1*** | A804D^1^ | This study. |
| ***DHR1*** | E831G^2^ | (3) |
| ***DHR1*** | E831K^2^ | (3) |
| ***DHR1*** | F837L^2^ | (3) |
| ***DHR1*** | M857I^1^ | This study. |
| ***DHR1*** | E1037K^1^ | This study. |
| ***DHR1*** | E1037Q^1^ | This study. |

**Superscript**: Found as spontaneous suppressor (1) or by error-prone PCR (2).
